# Nuclear myosin VI maintains replication fork stability

**DOI:** 10.1101/2022.07.28.501567

**Authors:** Jie Shi, Kristine Hauschulte, Ivan Mikicic, Srijana Maharjan, Valerie Arz, Jan B. Heidelberger, Jonas V. Schaefer, Birgit Dreier, Andreas Plückthun, Petra Beli, Helle D. Ulrich, Hans-Peter Wollscheid

**Author notes:** Lead contact & correspondence and.

## Abstract

The actin cytoskeleton is of fundamental importance for cellular structure and plasticity. However, abundance and function of filamentous (F-) actin in the nucleus are still controversial. Here we show that the actin-based molecular motor myosin VI contributes to the stabilization of stalled or reversed replication forks. In response to DNA replication stress, myosin VI associates with stalled replication intermediates and cooperates with the AAA ATPase WRNIP1 in protecting these structures from DNA2- mediated nucleolytic attack. Using nuclear localization sequence (NLS) and ubiquitin E3-fusion DARPins to manipulate myosin VI levels in a compartment-specific manner, we provide evidence for the direct involvement of myosin VI in the nucleus and against a contribution of the abundant cytoplasmic pool during the replication stress response.

## Main

Complete and correct duplication of the genome in each cell cycle is crucial for genome stability in proliferating cells. One of the many protective responses to DNA replication stress is the reversal of replication forks, involving a reannealing of the parental strands and a joining of the newly synthesized strands into a four-way Holliday junction-like structure^1, 2^. However, fork reversal, mediated by DNA-remodeling factors such as RAD51, SMARCAL1, HLTF and ZRANB3^3–5^, can also be detrimental for genome stability. Due to their structure resembling a one-ended double strand break (DSB), reversed forks can become targets of nucleolytic attack by nucleases such as DNA2 and MRE11, resulting in fork instability and collapse^6^.

The actin cytoskeleton exerts a fundamental role in cell mechanics, motility and intracellular transport. F-actin is highly abundant in the cytoplasm but barely detectable in the nucleus, where its functional relevance is still controversially discussed^7, 8^. Recent discoveries have connected nuclear F-actin to genome maintenance pathways such as DSB repair, DNA replication and maintenance of nuclear architecture^9–12^. If and how myosins in their function as actin-based molecular motor proteins participate in these processes is still poorly understood. The myosin superfamily comprises more than 20 distinct classes in humans, of which only a few have been shown to exert nuclear functions^13^.

Myosin VI, the only minus end-directed myosin characterized to date^14^, is well known for its contribution to multiple steps of the transcriptional process^15–17^. We recently identified a region adjacent to its C-terminal cargo-binding domain as a ubiquitin-interacting domain (MyUb, Fig. 1a)^18^. Pulldown assays with a GST-MyUb construct, followed by SILAC-based quantitative mass spectrometry (Fig. 1b), identified 490 proteins with an at least 2-fold enrichment over the GST control (FDR>0.05), including 346 proteins annotated with the gene ontology (GO) cellular compartment “nucleus”. In line with its known function, GO term analysis of the MyUb interactome showed transcription-associated proteins as the most prominently enriched category (Fig. 1c). In addition, we found many DNA replication-associated factors, suggesting a yet unidentified function of myosin VI at the replisome (Fig. 1d). Immunoprecipitation (IP) experiments upon overexpression of GFP-tagged proteins (Fig. S1b) or GST-pulldown experiments followed by immunoblotting with antibodies against endogenous proteins (Fig. 1e) validated many of the candidates identified in our proteomic screen as genuine interaction partners of myosin VI.

**Fig. 1.**
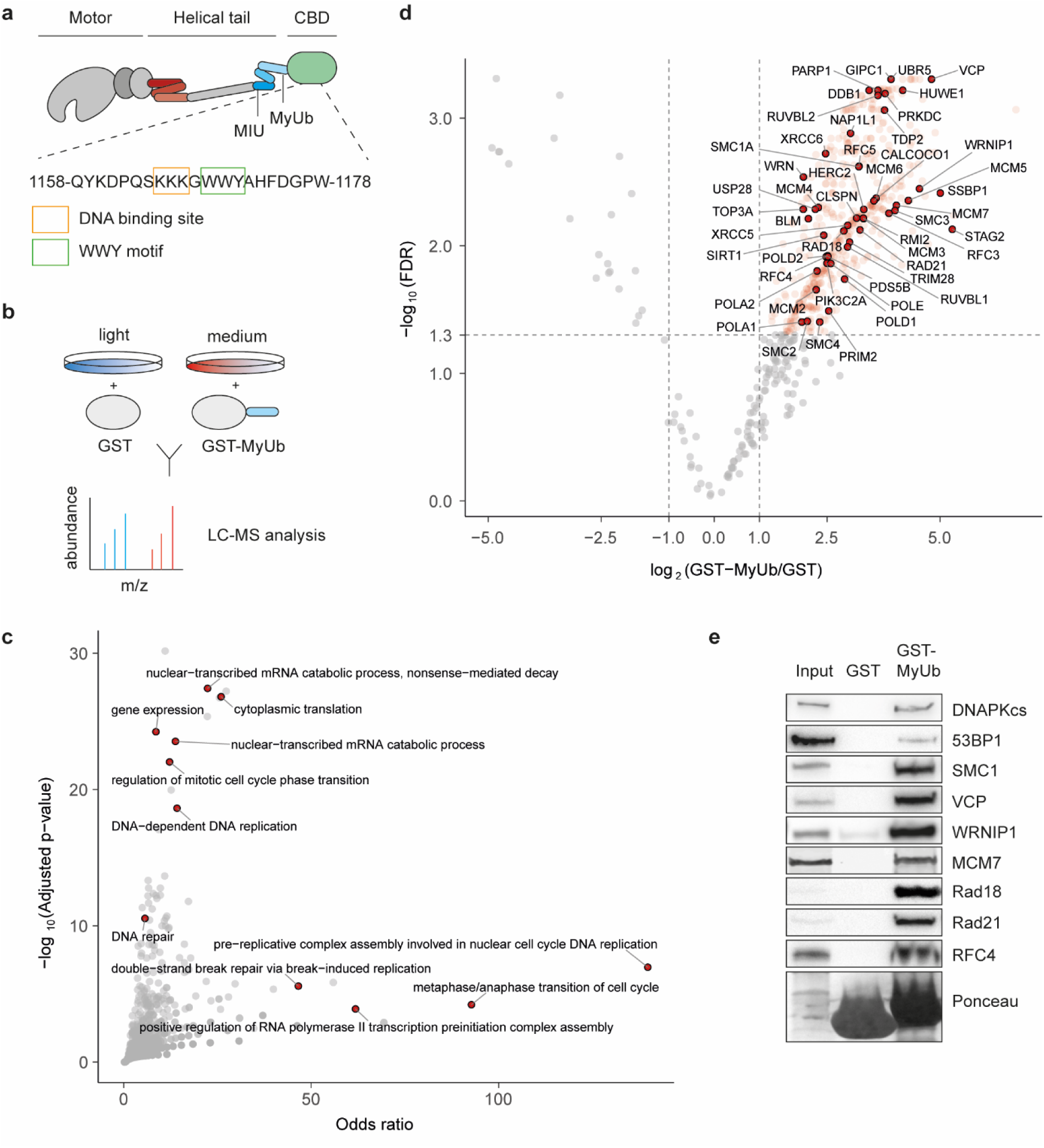
Myosin VI interacts with the replisome. **a**, Schematic representation of myosin VI (adapted from Magistrati and Polo ^39^) showing the position of the ubiquitin binding domains MIU and MyUb domain (blue) adjacent to the cargo-binding domain (CBD, green). Three-helix bundle at the N-terminal tail is indicated in red. The amino acid sequence shows a triple-Lys repeat involved in DNA binding^16^ (orange box) and the WWY motif (green box), a well-characterized protein interaction site. Amino acid numbering is according to the short isoform (isoform 2). **b**, Set-up of the SILAC experiment for identification of MyUb interaction partners. **c**, GO term analysis (GO biological process) of proteins identified to interact with the MyUb domain (fold change > 4, FDR < 0.05) using EnrichR. **d**, Volcano plot of protein groups identified in the SILAC interactome experiment. Mean log2 fold change of all replicates between GST-MyUb and GST are plotted against the −log10 FDR. Significantly enriched proteins are shown in red (fold change > 2, FDR < 0.05). Interactors involved in DNA replication and repair are highlighted and labeled. **e**, Validation of selected candidates by pulldown assays from total cell lysates with recombinant GST- MyUb, followed by western blotting with antibodies against endogenous proteins as indicated. Ponceau S staining shows equal loading of GST and GST-MyUb.

To assess a potential role of myosin VI during DNA replication, we measured replication speed using DNA fiber assays, where nascent DNA is labeled consecutively with two thymidine analogues, CldU and IdU. Knockdown of myosin VI indeed led to a reduction in overall DNA replication speed, suggesting its requirement for efficient DNA replication (Fig. 2a).

**Fig. 2.**
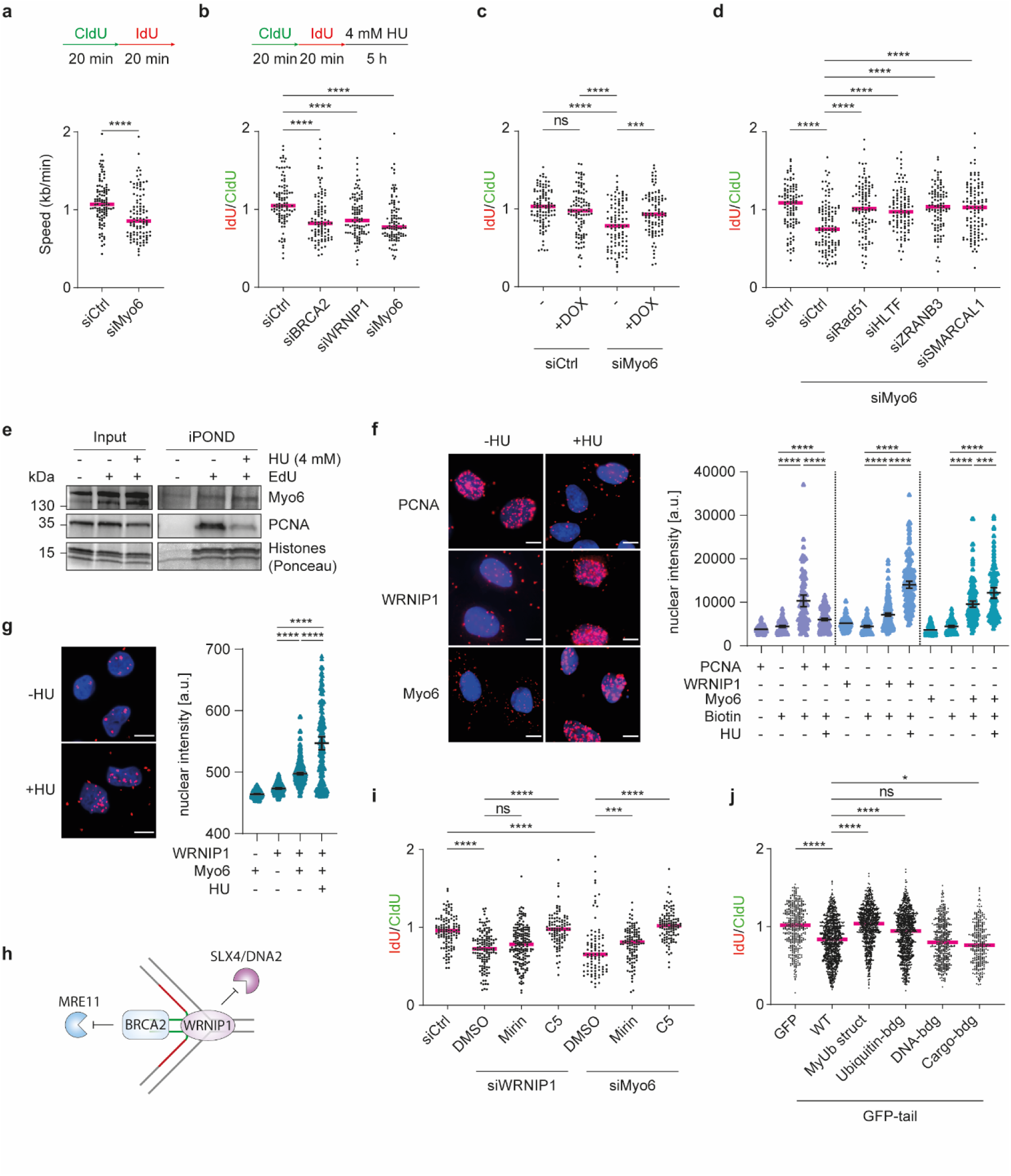
Myosin VI protects stalled replication forks from DNA2-mediated cleavage. **a**, Myosin VI is required for efficient unperturbed DNA replication. Top: Schematic representation of fiber assay conditions. Bottom: U2OS cells were transfected with siRNAs as indicated for 72 h, followed by fiber assays. Dot plots and median values of tract lengths. Replication speed was measured from total track length (CldU+IdU). Significance levels were calculated using the Mann-Whitney test from at least 100 fibers per sample (****: p<0.0001). **b**, Depletion of myosin VI via siRNA causes erosion of stalled replication forks in U2OS cells. Top: Schematic representation of fiber assay conditions. Bottom: U2OS cells were transfected with siRNAs as indicated for 72 h, followed by fiber assays. IdU/CldU ratios are shown as dot plots with median values. Significance levels were calculated using the Mann-Whitney test from at least 100 fibers per sample (ns: not significant, ****: p<0.0001, ***: p<0.001). **c**, GFP-myosin VI complements the loss of endogenous myosin VI. U2OS cells harboring DOX-inducible GFP-myosin VI were transfected with siRNAs for 72 h and treated with 20 ng/ml DOX for 24 h as indicated, followed by fiber assays performed as shown in panel **b**. **d**, Myosin VI-dependent fork protection requires fork reversal. U2OS cells were transfected with siRNAs for 72 h as indicated, followed by fiber assays performed as shown in panel **b**. **e**, iPOND assays show localization of myosin VI at replication forks. U2OS cells were pulsed with EdU for 30 min, followed by a subsequent 5 h treatment with 4 mM HU where indicated. Proteins associated with newly synthesized DNA were isolated using a standard iPOND protocol and visualized using western blotting with antibodies against PCNA and myosin VI. To control efficient chromatin isolation, Histones were visualized via Ponceau S staining. **f**, SIRF assays confirm the presence of myosin VI at replication forks. U2OS cells were pulsed with EdU for 30 min, followed by a subsequent 5 h treatment with 4 mM HU where indicated. Following the click-IT reaction with biotin-azide, PLA was performed using a biotin-specific antibody in combination with the indicated antibodies. Left: representative images with Hoechst staining in blue and PLA signals in magenta (scale bar = 10 µm). Right: dot plots of PLA signal intensities with mean values -/+ 95% confidence intervals. Significance levels were calculated using the Mann-Whitney test from at least 100 nuclei per sample (****: p<0.0001, ***: p<0.001). **g**, Enhanced interaction of myosin VI with WRNIP1 upon replication stress. U2OS cells were treated for 5 h with 4 mM HU where indicated, followed by a standard PLA. Left: representative images with Hoechst staining in blue and PLA signals in magenta (scale bar = 10 µm). Right: dot plots of PLA signal intensities with mean values -/+ 95% confidence intervals. Significance levels were calculated using the Mann-Whitney test from at least 100 nuclei per sample (****: p<0.0001). **h**, Schematic representation of different fork protection mechanisms by WRNIP1 and BRCA2 according to Porebski et al.^19^. **i**, Inhibition of DNA2 restores fork stability in WRNIP1- and myosin VI- deficient cells. U2OS cells were transfected with siRNAs for 72 h and treated with nuclease inhibitors (mirin and C5 for MRE11 and DNA2, respectively) or DMSO for 5 h as indicated, followed by fiber assays performed as shown in panel **b**. **j**, Motor- and MyUb-domains of myosin VI are required for its function in fork protection. GFP-tail *wildtype* (WT) and mutants were overexpressed in U2OS cells for 48 h as indicated, followed by fiber assays performed as shown in panel **b**. Combined data from at least 3 independent replicates are shown. Detailed information about the respective mutations are given in Fig. **S2f**. For **a, b, c, d, e, f, g, i:** A representative experiment from three independent replicates is shown. For **a, b, c, d, i, j:** Knockdown efficiencies and overexpression levels are shown in Fig. **S2**.

The AAA ATPase WRNIP1 has been implicated in genome maintenance as a protector of reversed replication forks^19, 20^. Considering its identification as an interaction partner of myosin VI (Fig. 1d,e,S1b), we asked whether the replication problems upon myosin VI depletion were linked to a defect in fork protection. To this end, we labeled cells with CldU and IdU for 20 min each, followed by a 5 h treatment with hydroxyurea (HU) to stall replication forks (Fig. 2b). As degradation of newly replicated DNA leads to a shortening of the second (IdU) tract, analysis of the IdU/CldU ratio allows an estimation of the extent of fork degradation. According to their well-established roles as replication fork protectors, siRNA-mediated depletion of WRNIP1 and BRCA2^21^ resulted in nascent strand degradation as indicated by a reduction in the IdU/CldU ratio (Fig. 2b). Notably, myosin VI depletion reduced the IdU/CldU ratio to a similar extent, suggesting that myosin VI is essential for preventing nuclease-mediated degradation of reversed forks (Fig. 2b). To exclude off-target effects, we carried out rescue experiments using a cell line expressing siRNA-resistant GFP-myosin VI under the control of a doxycycline (DOX) -inducible promoter (Fig. S2c). In control cells expressing endogenous myosin VI, addition of DOX did not significantly alter the stability of stalled replication forks (Fig. 2c, lanes 1 and 2). However, in myosin VI-depleted cells, we observed a rescue of fork protection upon DOX-induced restoration of myosin VI levels (Fig. 2c, lanes 3 and 4), thus verifying the direct correlation between replication fork stability and myosin VI abundance. Furthermore, co-depletion of the fork remodelers RAD51, HLTF, SMARCAL1 or ZRANB3 together with myosin VI completely abolished nascent strand degradation (Fig. 2d), indicating that the defect in fork stability induced by myosin VI depletion depends on the prior action of the fork remodelers. Thus, myosin VI appears to protect reversed replication forks, but it does not prevent fork reversal.

In contrast to other fork protectors, myosin VI primarily localizes to the cytoplasm and its abundance in the nucleus – like that of F-actin – is low. To investigate a potential physical association of myosin VI with ongoing and stalled or reversed replication forks, we therefore utilized iPOND (isolation of proteins on nascent DNA) with western blotting to focus specifically on chromatin-associated factors^22^. PCNA is known to dissociate from newly replicated DNA upon replication stress^23^, and this pattern was reproducible in our hands (Fig. 2e). In agreement with the observed interactions of myosin VI with replisome components (Fig. 1c, e), we detected myosin VI at unperturbed replication forks (Fig. 2e). Unlike PCNA, however, myosin VI association was not diminished upon HU treatment. To achieve a more quantitative assessment, we used SIRF (*in situ* protein interaction with nascent DNA replication forks) assays, which detect the co-localization of a protein of interest with nascent, EdU-labeled DNA via proximity ligation^24^. Again, the PCNA signal was lost under conditions of replication stress, while both myosin VI and WRNIP1 showed enhanced association with EdU-positive nascent DNA upon HU treatment (Fig. 2f), suggesting an enrichment of both myosin VI and WRNIP1 at stalled forks.

Having established the interaction of WRNIP1 with the MyUb domain of myosin VI (Fig. 1e, S1b), we utilized proximity ligation assays (PLA) to validate this interaction in living cells using antibodies against the endogenous proteins (Fig. 2g). Strikingly, the PLA signal was prominently enhanced under conditions of replication stress, suggesting that the proteins preferentially interact at stalled replication forks (Fig. 2g). Unlike BRCA2, which is thought to protect the ends of the regressed arm from MRE11-dependent degradation^21^, WRNIP1 was reported to prevent attack by SLX4/DNA2 at the four-way junction^19^ (Fig. 2h). To specify the nature of myosin VI activity at reversed forks, we performed DNA fiber assays in the presence of the MRE11- or DNA2-specific inhibitors mirin or C5, respectively. Consistent with previous findings, mirin treatment did not rescue nascent strand degradation in WRNIP1-depleted cells, while DNA2 inhibition led to a full stabilization of reversed forks (Fig. 2i)^19^. Use of the inhibitors in myosin VI-depleted cells resulted in a very similar pattern (Fig. 2i), suggesting that myosin VI cooperates with WRNIP1 to protect reversed replication forks from DNA2- mediated nucleolytic attack.

Next, we investigated the molecular characteristics of myosin VI-dependent fork protection. Overexpression of a motor-deficient variant (“GFP-tail”) resulted in nascent strand degradation similar to myosin VI depletion (Fig. 2j), demonstrating the importance of its motor activity for fork protection. By exploiting this dominant-negative effect, we addressed the contributions of multiple interaction sites of myosin VI (Fig. S2f) to the replication stress response. It was previously shown that mutation of the RRL motif within the MyUb domain leads to destabilization of its helical structure^18^. In line with the multitude of replication factors that interact with this domain, mutation of the RRL motif to AAA abolished the dominant-negative effect of the GFP-tail construct (Fig. 2j, lane3). Whereas a combination of point mutants in the MIU (A1013G)^25^ and MyUb (I1072A)^18^ domains revealed a contribution of ubiquitin binding to myosin VÍs activity in fork protection, its DNA^16^- and WWY^26^- mediated cargo-binding activities (Fig. 1a,S2f) seem to be less important (Fig. 2j).

Actin filaments are of a transient nature and difficult to detect in the nucleus because of their high cytoplasmic abundance. An actin-specific nanobody fused to a nuclear localization signal (NLS), termed nuclear actin chromobody (nAC), has proven to be a valuable instrument in visualizing nuclear F-actin specifically^27^. However, manipulation of nuclear F-actin remains challenging due to the involvement of monomeric actin in chromatin remodeling complexes^28^ and its association with RNA polymerase complexes^29–31^. Inspired by the nAC technology, we aimed to develop tools to manipulate the stability and localization of endogenous myosin VI. To obtain a myosin VI-specific affinity probe, we employed a ribosome display library of designed ankyrin repeat proteins (DARPins), which consist of stacked repeat modules with a randomized surface. They can be selected to bind proteins with antibody-like selectivity and affinity^32–34^. Unlike antibodies, DARPins fold under the reducing conditions of the cytoplasm and the nucleus and can thus be expressed in these compartments. Using a biotinylated tail fragment of myosin VI (aa 992 - 1031) as bait, we obtained 54 candidates from the library that were further screened in GST-pulldown experiments (Fig. S3a). Five of these clones were further tested for their ability to deplete endogenous myosin VI from cellular lysates, using a non-selective DARPin (E3_5)^34^ as negative control. One clone, “M6G4”, effectively depleted myosin VI from the lysate and was therefore selected as the target-binding module for the myosin VI-specific tools (Fig. 3a, S3b).

**Fig. 3.**
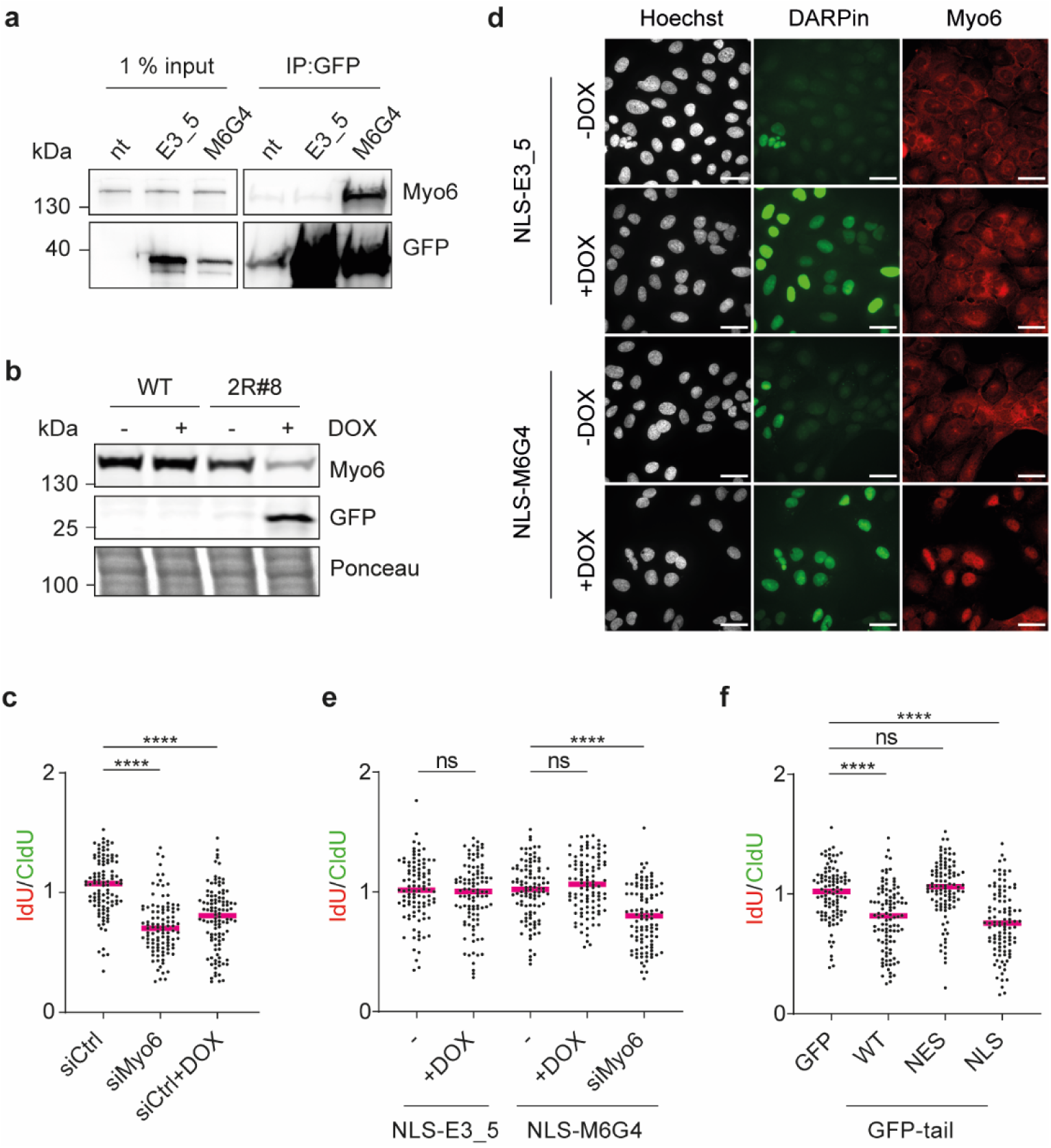
Nuclear but not cytoplasmic myosin VI is active in fork protection. **a**, DARPin M6G4 depletes myosin VI from cellular lysates. HEK293T cells were transfected with GFP- E3_5 control DARPin or GFP_M6G4 anti-myosin VI DARPin for 24 h as indicated. Immunoprecipitations (IPs) against GFP using GFP-trap beads (Chromotek) were performed, followed by western blotting with antibodies against myosin VI and GFP. **b**, DOX-induced degradation of myosin VI via a DARPin-based construct. A U2OS Flp-In T-REx single-cell clone harboring a DOX-inducible GFP-M6G4-2RING fusion construct (2R#8) was treated with 20 ng/ml DOX for 24 h. Cellular lysates were analyzed via western blotting using antibodies against myosin VI and GFP. Ponceau S staining was used to compare loading between samples. **c**, DARPin-2RING fusion-mediated degradation of myosin VI confirms its contribution to fork protection. U2OS Flp-In T-REx cells harboring DOX-inducible GFP-M6G4-2RING (2R#8) were transfected with the siRNA-pool against myosin VI for 72 h and treated with DOX for 24 h as indicated, followed by fiber assays performed as shown in Fig. 2b. **d**, DARPin-mediated re-localization of myosin VI to the nucleus. U2OS Flp-In T-REx cells harboring DOX-inducible GFP-M6G4-NLS or GFP-E3_5-NLS (control) were treated with 20 ng/ml DOX for 24 h where indicated. Subsequently, immunofluorescence analyses were performed using myosin VI-specific antibodies (red channel). DARPin expression and localization were monitored in the GFP channel. Nuclei were stained with Hoechst (white signal) (scale bar = 40 µm). **e**, Depletion of cytoplasmic myosin VI has no effect on fork stability. U2OS Flp-In T-REx cells harboring DOX-inducible GFP-M6G4-NLS or GFP-E3_5-NLS (control) were transfected with siRNA#10 against myosin VI and treated with DOX as indicated, followed by fiber assays performed as shown in Fig. 2b. **f**, Inhibition of nuclear but not cytoplasmic myosin VI leads to fork de-stabilization upon replication stress. U2OS cells were transfected with compartment-specific GFP-tail constructs for 48 h as indicated, followed by fiber assays performed as shown in Fig. 2b. For **c, e, f:** A representative experiment from three independent replicates is shown. Knockdown efficiencies and overexpression levels are shown in Fig. **S3**.

Next, we adapted a recently published inducible degradation system based on the ubiquitin protein ligase RNF4^35^ (Fig. S3c). A fusion construct of DARPin M6G4 with two RING finger domains of RNF4 (M6G4-2RING) was stably integrated in cells under the control of a DOX-inducible promoter. A single-cell clone termed 2R#8 showed efficient degradation of endogenous myosin VI in a time- and DOX-dependent manner (Fig. 3b, S3d,e). Importantly, depletion of myosin VI via M6G4-2RING resulted in a destabilization of stalled forks, comparable to siRNA-mediated myosin VI depletion (Fig. 3c), providing additional support for the specificity of the phenotype.

Having verified the selectivity of the M6G4 probe, we asked whether fork stability was regulated by the nuclear or the cytoplasmic pool of myosin VI. We found that inducible expression of a GFP-tagged fusion construct of M6G4 to a 3 x NLS resulted in a nearly complete localization of myosin VI to the nuclear compartment (Fig. 3d), while the analogous GFP-NLS-E3_5 control construct did not afford significant changes in the subcellular distribution. Fiber assays in cells expressing either the myosin VI- specific- or the control- NLS-DARPin did not show significant degradation of newly replicated DNA (Fig. 3e), suggesting that depletion of cytoplasmic myosin VI has little or no influence on fork stability. Unfortunately, our attempts to selectively deplete myosin VI from the nucleus by fusion of an analogous nuclear export signal (NES) was deemed impracticable since the NES-DARPin fusion, synthesized in the cytosol, probably does not enter the nucleus, and an NES-anti-GFP DARPin used as a test case, could thus not export GFP (Fig. S3h).

As an alternative approach, we therefore expressed motor-deficient myosin VI mutants (“NLS/NES- tail”) intended as dominant-negative alleles that would compete with endogenous myosin VI for functional interactions in the respective subcellular compartments. Whereas expression of nuclear NLS-tail caused significant degradation of nascent DNA, expression of cytoplasmic NES-tail had no effect (Fig. 3f), strongly suggesting that the compartment relevant for myosin VI activity in fork protection is the nucleus rather than the cytoplasm.

The requirement of myosin VÍs motor domain for its function in fork protection implied a mobility-dependent mechanism (Fig. 2j). This might involve an active transport of fork-protecting factors such as WRNIP1 towards stalled or reversed forks (Fig. 4a) or, alternatively, a transport of fork-destabilizing factors such as pertinent nucleases away from the sites of fork stalling (Fig. 4b). To differentiate between these models, we used SIRF to test whether myosin VI affected the recruitment of WRNIP1 to unperturbed or stalled replication forks. Consistent with our previous results (Fig. 2f), control cells expressing myosin VI afforded a WRNIP1 signal at unperturbed forks that increased after HU treatment (Fig. 4c). Knockdown of myosin VI did not significantly affect association of PCNA with replication forks (Fig. 4c, left panel) and WRNIP1 recruitment to unperturbed replication forks. However, under conditions of replication stress, we scored a clear defect in WRNIP1 accumulation at forks upon depletion of myosin VI, arguing for a model where myosin VI positively regulates WRNIP1’s association with reversed replication forks (Fig. 4a). Conversely, WRNIP1 depletion did not affect localization of myosin VI to replication forks (Fig. S4).

**Fig. 4.**
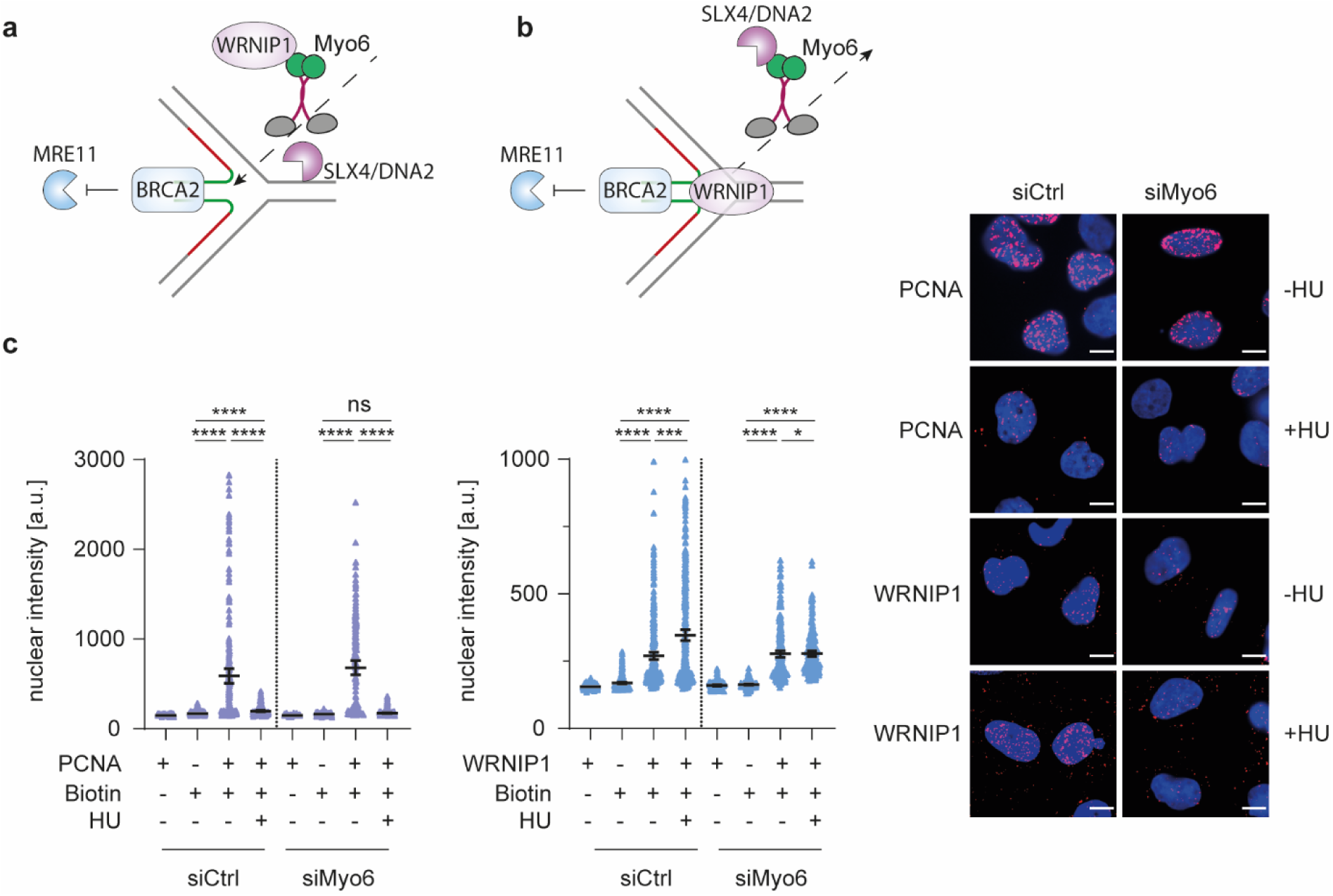
Myosin VI is required for efficient localization of WRNIP1 to stalled forks. **a, b**, Models of how myosin VI could mediate fork protection in its role as a motor protein. **c**, Myosin VI depletion interferes with efficient fork localization of WRNIP1. U2OS cells were transfected with siRNAs for 72 h as indicated, followed by SIRF assays. Left: dot plots of PLA signal intensities with mean values -/+ 95% confidence intervals. Significance levels were calculated using the Mann-Whitney test from at least 100 nuclei per sample (ns, non-significant, ****: p<0.0001, ***: p<0.001, *: p<0.05). Right: representative images with Hoechst staining in blue and PLA signals in magenta (scale bar = 10 µM). A representative experiment from three independent replicates is shown. Knockdown efficiency is shown in Fig. **S4**. Supplemental information

Our findings connect the actin-based motor protein myosin VI to a defined pathway of replication fork protection that maintains genome stability under conditions of replication stress. Using an unbiased mass spectrometry approach in combination with *in situ* localization studies, we found myosin VI to accumulate at stalled replication forks in response to nucleotide depletion, and functional assays have revealed a contribution to the protection of reversed forks from nucleolytic attack by DNA2. A scenario where myosin VI acts by mediating the transport of the fork protection factor WRNIP1 to its sites of action is consistent with our protein interaction and localization data (Fig. 4c). The notion that the motor domain of myosin VI is required for its function suggests a role in shuttling; however, as myosin VI has also been shown to act in an anchoring fashion^36^, we cannot exclude a model where myosin VI stabilizes the fork protection complex at the junction between parental and reversed strands in a static manner. Likewise, although the identification of WRNIP1 as an interaction partner of myosin VI by mass spectrometry suggests a direct physical effect of the motor protein on the recruitment of the ATPase, we cannot exclude indirect interactions between the two proteins. On the side of myosin VI, we found the ubiquitin-binding MyUb domain to be functionally important. Intriguingly, WRNIP1 also harbors a well-characterized ubiquitin-binding (UBZ) domain; interactions relevant for fork protection might therefore be mediated by a common association of both factors with ubiquitylated structures. In contrast, DNA binding by myosin VI does not appear to be important in this context, as the relevant mutant did not cause any fork destabilization.

Beyond the functional interaction of myosin VI and WRNIP1, our data support and expand recent evidence for nuclear functions of the actin cytoskeleton in genome maintenance. Although we did not directly address nuclear F-actin, the requirement of the myosin VI motor domain for fork protection (Fig. 2j,3f) strongly suggests a mechanism based on the movement of the myosin on nuclear actin filaments rather than invoking an actin-independent mechanism. However, WASp, a positive regulator of ARP2/3 dependent actin-polymerization, was recently shown to modulate RPA-regulated signaling upon genotoxic insult^37^. The authors convincingly demonstrate an actin-independent role of WASp as “chaperone” for RPÁs ssDNA binding. Likewise, we cannot rule out additional, actin-independent functions for myosin VI.

Finally, while technical limitations have so far precluded firm evidence against an influence of the cytoplasmic actin cytoskeleton on genome maintenance, our newly designed tools in combination with classical dominant-negative approaches have provided clear evidence for the relevance of the nuclear pool of myosin VI for fork protection while excluding myosin VI-related cytoplasmic signaling events. The formation of actin filaments inside the nucleus upon replication stress, detected by Lamm et al.^38^ raises speculations about the relevance of the unique minus-end directionality of myosin VI and the possible orientation of actin filaments forming in the vicinity of reversed forks. We also envision the involvement of other myosins, e.g. myosin I or myosin V^10^, opening the possibility for a competition between minus- and plus-end-directed motors. Probing the role of other myosins as well as actin cytoskeleton proteins such as bundling, capping, assembly or disassembly factors will thus be important for future studies. Taken together, our discovery of the requirement of myosin VI-dependent transport or tethering for the protection of reversed replication forks, possibly controlled by ubiquitin binding, adds to the accumulating evidence for a key role of nuclear actin filaments in genome maintenance and paves the way for exploring new layers of regulation of nuclear transactions by a set of proteins better known for their role in cytoplasmic signaling.

## Methods

### Cell lines, cultivation and treatments

U2OS, HeLa and HEK293T cells were maintained in DMEM containing 10% fetal bovine serum, L-glutamine (2 mM), penicillin (100 U/ml), and streptomycin (100 µg/ml) (Thermo Fisher Scientific). U2OS Flp-In T-REx cell lines were maintained in DMEM containing 10% fetal bovine serum, L-glutamine, penicillin, streptomycin and blasticidin (5 µg/ml) (Invivogen). All cell lines were cultured in humidified incubators at 37°C with 5% CO_2_. Treatments were performed with hydroxyurea (4 mM, Merck), the MRE11 inhibitor mirin (25 µM, Merck) or the DNA2-specific inhibitor C5 (25 µM, AOBIOUS) for 5 h.

For SILAC labeling, HeLa cells were cultured for at least 5 passages in DMEM containing either L-arginine and L-lysine (Merck) or L-arginine [^13^C6] and L-lysine [^2^H4] (Cambridge Isotope Laboratories).

### Transfections

For overexpression purposes, HEK293T were transfected using polyethyleneimine (PEI) (Polysciences), other cell types were transfected using Fugene HD (Promega) or Lipofectamine 2000 (Life technologies) according to the manufactureŕs instructions. All expression constructs used in this study are listed in Supplementary Table 1.

For knockdowns, cells were transfected with siRNAs using Lipofectamine RNAiMAX (Life Technologies) according to the manufactureŕs instructions at a final RNA concentration of 20 nM for 72 h. Knockdown of myosin VI was achieved with a pool of 4 different siRNAs (Hs_MYO6_5 FlexiTube siRNA, Hs_MYO6_7 FlexiTube siRNA, Hs_MYO6_8 FlexiTube siRNA and Hs_MYO6_10 FlexiTube, Qiagen). For rescue experiments in the U2OS Flp-In cell line expressing GFP-myosin VI, a single siRNA targeting the 3’-UTR of the myosin VI transcript (Hs_MYO6_10 FlexiTube siRNA) was used. RAD51 and ZRANB3 knockdowns were performed using a pool of two independent siRNAs each. A list of all siRNAs used in this study can be found in Supplementary Table 3.

### Generation of stable cell lines

U2OS Flp-In T-REx cell lines for DOX-inducible expression were generated by co-transfection of the respective pDEST-FRT-TO construct with the pOG44 Flp-Recombinase (Supplementary Table 1). 24 h post-transfection, cells were selected with 100 µg/ml hygromycin (Invivogen) for 10 days. Hygromycin-resistant cells were sorted for GFP-positive clones using a BD FACS Aria III SORP instrument. Single-cell clones were tested for construct expression and myosin VI depletion after DOX treatment by western blotting using GFP- and myosin VI-specific antibodies (Supplementary Table 2).

### Generation of plasmids

Fragments were inserted via restriction/ligation cloning or following PCR amplification with specific oligonucleotides, listed in Supplementary Table 4. For Gateway cloning, Gateway® LR Clonase® II enzyme mix (Thermo Fisher Scientific) was used according to the manufactureŕs instructions. Detailed information about individual constructs will be provided upon request.

### Site-directed mutagenesis

Site-directed mutagenesis was performed using Pfu Turbo DNA Polymerase (Agilent). The amplification product was digested with DpnI (New England Biolabs), *E. coli* TOP10 cells were transformed with the construct followed by sequence verification. Oligonucleotides for mutagenesis are listed in Supplementary Table 4.

### Protein production and purification

GST fusion proteins were produced in *E. coli* Bl21 (DE3) cells at 37°C for 4 h after induction with 1 mM IPTG (Generon) at an OD_600_ of 0.8. Cells were pelleted and lysed by sonication in PBS/0.1% Triton X- 100 (Merck) supplemented with protease inhibitor cocktail (SIGMAFAST). Clarified supernatants were incubated with 1 ml of GSH-Sepharose beads (Cytiva) per liter of bacterial culture. After 2 h at 4°C, the beads were washed with PBS/0.1% Triton X-100 and maintained in storage buffer (50 mM Tris, pH 7.4, 100 mM NaCl, 1 mM EDTA, 1 mM DTT, and 10% glycerol).

Expression of DARPins with N-terminal MRGS(H)_8_ tag and myosin VI (aa 992-1031) with N-terminal MRGS(H)_8_ and C-terminal Avi tag in *E. coli* BL21 (DE3) was induced with 1 mM IPTG for 20 h at 18°C. Cells were resuspended in buffer A (50 mM Tris-HCl pH 7.4, 250 mM NaCl, 10% glycerol, 1 mM DTT, 20 mM imidazole) and lysed by sonication. The clarified supernatant was subjected to affinity chromatography on Ni-NTA resin (Qiagen), and eluted protein was rebuffered using PD 10 columns (Thermo Fisher Scientific) in storage buffer (50 mM Tris, pH 7.4, 100 mM NaCl, 1 mM EDTA, 1 mM DTT, and 10% glycerol).

Myosin VI (aa 992-1031) with N-terminal MRGS(H)_8_ and C-terminal Avi tag was biotinylated *in vivo* by co-expressing biotin-ligase BirA (pBirAcm from Avidity) in *E.coli* BL21 (DE3). 50 µM biotin was added to the growth medium (LB) before induction with IPTG.

### GST-pulldown assay coupled to mass spectrometry

For SILAC experiments, 8x10^7^ HeLa cells were lysed in 2 ml JS buffer (100 mM HEPES pH 7.5, 50 mM NaCl, 5% glycerol, 1% Triton X-100, 2 mM MgCl_2_, 5 mM EGTA, 1 mM DTT), supplemented with protease inhibitor cocktail (SIGMAFAST) and Benzonase® (Merck). 50 µg of GST and 70 µg of GST-MyUb fusion protein immobilized on 50 µl GSH-Sepharose beads were incubated with 1 ml of cellular lysate for 2 h at 4°C. Beads were washed 5 times in 1 ml JS buffer. Labels were switched in 2 out of 4 biological replicates. SILAC samples were pooled during the last wash. Bound proteins were eluted in 2x NuPAGE LDS Sample Buffer (Life Technologies) supplemented with 1 mM dithiothreitol, heated at 70 °C for 10 min, alkylated by addition of 5.5 mM chloroacetamide for 30 min, and separated by SDS–PAGE on a 4–12% gradient Bis–Tris gel (Invitrogen). Proteins were stained using the Colloidal Blue Staining Kit (Life Technologies) and digested in-gel using trypsin (Serva). Peptides were extracted from the gel and desalted using reversed-phase C18 StageTips.

Peptide fractions were analyzed on a quadrupole Orbitrap mass spectrometer (Q Exactive Plus, Thermo Fisher Scientific) equipped with a UHPLC system (EASY-nLC 1000, Thermo Fisher Scientific). Peptide samples were loaded onto C18 reversed-phase columns (25 cm length, 75 μm inner diameter, 1.9 μm bead size, packed in-house) and eluted with a linear gradient from 1.6 to 52% acetonitrile containing 0.1% formic acid in 90 min. The mass spectrometer was operated in a data-dependent mode, automatically switching between MS and MS2 acquisition. Survey full scan MS spectra (m/z 300–1,650, resolution: 70,000, target value: 3e6, maximum injection time: 20 ms) were acquired in the Orbitrap. The 10 most intense ions were sequentially isolated, fragmented by higher energy C-trap dissociation (HCD) and scanned in the Orbitrap mass analyzer (resolution: 35,000, target value: 1e5, maximum injection time: 120 ms, isolation window: 2.6 m/z). Precursor ions with unassigned charge states, as well as with charge states of +1 or higher than +7, were excluded from fragmentation. Precursor ions already selected for fragmentation were dynamically excluded for 20 s.

Raw data files were analyzed using MaxQuant (version 1.5.2.8)^40^. Parent ion and MS2 spectra were searched against a reference proteome database containing human protein sequences obtained from UniProtKB (HUMAN_2016_05) using the Andromeda search engine^41^. Spectra were searched with a mass tolerance of 4.5 ppm in MS mode, 20 ppm in HCD MS2 mode, strict trypsin specificity, and allowing up to two mis-cleavages. Cysteine carbamidomethylation was searched as a fixed modification, whereas protein N-terminal acetylation, methionine oxidation, GlyGly (K), and N-ethylmaleimide modification of cysteines (mass difference to cysteine carbamidomethylation) were searched as variable modifications. The Re-quantify option was turned on. The dataset was filtered based on posterior error probability (PEP) to arrive at a false discovery rate of below 1%, estimated using a target-decoy approach^42^. Statistical analysis and MS data visualization were performed using the R software environment (version 1.3.1093). Potential contaminants, reverse hits, hits only identified by site and hits with no unique peptides were excluded from the analysis. Statistical significance was calculated using a moderated t-test (limma package)^43^. GO term analysis (biological process) was performed using EnrichR^44^. Visualized GO terms were selected based on adjusted p-value, odds ratio and semantic uniqueness. To determine the number of nuclear proteins among MyUb interactors (fold change > 2, FDR < 0.05), GO cellular component annotations were retrieved from the STRING network tool^45^.

### GST-pulldown assays

For validation purposes, GST-pulldown assays were performed with lysates from 5x10^6^ unlabeled HeLa cells, and interactors were detected by western blotting using antibodies against endogenous proteins. To identify DARPins suitable in pulldown assays, screening was performed by incubating 10 µg GST (as control) or 14 µg GST-MyUb immobilized on 20 µl GSH-Sepharose beads with a final DARPin concentration of 1 µM in 200 µl PBS/0.1% Triton X-100. Beads were washed 3 times in 1 ml PBS/0.1% Triton X-100, boiled for 10 min in NuPAGE^®^ LDS Sample Buffer and subjected to SDS-PAGE. Detection was performed using Instant Blue protein stain (Biozol).

### Immunoprecipitation of GFP-tagged proteins

HEK293T cells were PEI-transfected with the respective plasmid (Supplementary Table 1) for 24 h, followed by lysis in JS buffer (100 mM HEPES pH 7.5, 50 mM NaCl, 5% glycerol, 1% Triton X-100, 2 mM MgCl_2_, 5 mM EGTA, 1 mM DTT) supplemented with protease inhibitor cocktail (SIGMAFAST) and Benzonase®. Cell lysates were cleared by centrifugation for 30 min at 4°C and incubated with GFP-trap magnetic agarose beads (Chromotek) for 1 h at 4°C. After 3 washes with JS buffer, beads were boiled for 10 min in NuPAGE^®^ LDS Sample Buffer and subjected to western blotting.

### iPOND

U2OS cells were labeled with 10 μM EdU (Merck) for 30 min. Subsequently, cells were fixed with 1% formaldehyde (Merck) for 10 min, followed by quenching with 125 mM glycine (Merck) for 10 min. After 2 washing steps with PBS/1% BSA, cells were collected by scraping, followed by permeabilization in 0.1% Triton X-100/PBS. Subsequently, cells were washed with 1% BSA/PBS and subjected to the Click-iT reaction in a solution containing 10 mM sodium ascorbate (Merck), 0.1 mM azide-PEG_3_-biotin conjugate (Merck) and 2 mM copper sulfate (Merck) for 30 min at room temperature. Cells were then washed twice in 1% BSA/PBS, lysed in 10 mM Tris-HCl pH 8.0, 140 mM NaCl, 1% Triton X-100, 0.1% sodium deoxycholate, 0.1% SDS, supplemented with SIGMAFAST protease inhibitor cocktail, and sonicated using a Bioruptor (Diagenode). Lysates were cleared by centrifugation for 45 min at 4°C in a table-top centrifuge and subjected to streptavidin-agarose beads (Thermo Fisher Scientific) overnight at 4°C. On the next day, beads were washed five times in 1% BSA/PBS and de-crosslinking was carried out for 30 min in NuPAGE^®^ LDS Sample Buffer at 95°C. For protein detection, samples were subjected to SDS-PAGE and immunoblotted with relevant antibodies (Supplementary Table 2).

### Immunofluorescence

For immunofluorescence analysis, cells were fixed with 4% paraformaldehyde (Merck) for 10 min and permeabilized for 5 min at room temperature with 0.1% Triton X-100. Subsequently, cells were incubated with primary antibodies for 1 h (α-myosin VI α-rabbit in a 1:400 dilution), followed by 3 x 5 min washing steps with PBS/0,1% Triton X-100 and incubation with secondary antibodies for 30 min at room temperature. Coverslips were mounted with ProLong™ Diamond Antifade Mountant (Thermo Fisher Scientific). Images were acquired with a Leica AF-7000 widefield microscope and analyzed with ImageJ.

### Immunoblotting

Samples were separated via SDS-PAGE and transferred to nitrocellulose membranes using the Trans-Blot Turbo® system (Bio Rad). Membranes were blocked for 1 h at room temperature in 5% milk/PBS/0.1% TWEEN-20 and incubated with primary antibodies (1:1000 dilution in PBS/0.1% TWEEN- 20/1% BSA) either for 1 h at room temperature or overnight at 4°C. Afterwards, membranes were washed with PBS/0.1% TWEEN-20 and incubated with secondary antibodies for 1 h at room temperature. Detection was performed by enhanced chemiluminescence using a Fusion FX (Vilbor lourmat) instrument after incubation with HRP-coupled secondary antibodies or by direct fluorescence using an Odyssey Clx imaging system (LI-COR) after incubation with secondary antibodies coupled to a fluorescent dye (Supplementary Table 2).

### Fiber assays

U2OS cells were labeled with 50 μM CldU (Merck) for 20 min and 50 μM IdU (Merck) for 20 min, respectively. Cells were trypsinized, resuspended in PBS and diluted to 1.75 x10^5^ cells/ml. Labeled cells were mixed with unlabeled cells at a ratio of 1:1. Lysis of the cells was carried out directly on microscopy slides, where 4 μl of the cells was mixed with 7.5 μl of lysis buffer (200 mM Tris-HCl pH 7.4, 50 mM EDTA, 0.5% SDS). After 9 min, the slides were tilted at an angle of 15-45° and the DNA fibers were stretched on the slides. The fibers were fixed in methanol/acetic acid (3:1) overnight at 4°C. Following fixation, the DNA fibers were denatured in 2.5 M HCl for 1 h, washed with PBS and blocked with 2% BSA/PBST for 40 min. The fibers were incubated with primary antibodies against CldU (Rat monoclonal anti-BrdU (clone BU1/75 (ICR1), Abcam) and IdU (Mouse monoclonal anti-BrdU (clone B44), BD Biosciences) (1:50 dilution) for 2.5 h, washed with PBST and incubated with secondary antibodies labeled with Alexa Fluor 488 and Alexa Fluor 647 (1:100 dilution). The slides were mounted in ProLong™ Diamond Antifade Mountant. Images of the DNA fibers were acquired using a Leica Thunder widefield microscope and analysis was carried out using Fiji ImageJ.

### Proximity Ligation Assays (PLA)

U2OS cells were seeded on coverslips with a confluency of 80%. Afterwards, cells were fixed in 4% paraformaldehyde for 10 min and permeabilized in 0.3% Triton X-100 for 10 min. PLA was then carried out using the Duolink® In Situ Red starter kit (Merck) according to the manufactureŕs instructions. Primary antibodies were used in a 1:100 dilution (α-WRNIP1 α-rabbit, α-Myosin VI α-mouse). In addition, Hoechst staining was included prior to mounting coverslips in ProLong™ Diamond Antifade Mountant. Images were acquired using a Leica Thunder widefield microscope and analysis was carried out using Fiji ImageJ.

### In situ analysis of protein interactions at DNA replication forks (SIRF)

For SIRF, cells were pulsed with 10 µM EdU for 10 min and then left untreated or treated with 4 mM HU for 5 h. After fixation in 4% paraformaldehyde for 10 min and permeabilization in 0.3% Triton X- 100 for 10 min, the Click-iT reaction was performed for 1 h at room temperature in PBS containing 2 mM copper sulfate, 10 µM azide-PEG_3_-biotin conjugate and 100 mM sodium ascorbate. PLA was then carried out as described above. Primary antibodies were used in a 1:100 (α-WRNIP1 α-rabbit, α- Myosin VI α-mouse, α-Biotin α-mouse) or 1:1000 dilution (α-PCNA, α-rabbit).

### DARPin selection and initial screening

To generate DARPin binders, biotinylated myosin VI (aa 992-1031) isoform 1 with N-terminal MRGS(H)_8_ and C-terminal Avi tag (^His^myosin VI (aa 992-1031)^Avi^) was immobilized on either MyOne T1 streptavidin-coated beads (Pierce) or Sera-Mag neutravidin-coated beads (GE Healthcare). The use of the type of beads was alternated during selection rounds. Ribosome display selections were performed essentially as described^46^, using a semi-automatic KingFisher Flex MTP 96 well platform.

The library includes N3C DARPins, displaying three internal and randomized ankyrin repeats as described earlier^34^. The initial C-cap was then replaced with a C-cap showing better stability towards unfolding implementing mutations in 5 amino acid positions^32, 47, 48^ to facilitate downstream experiments like protein fusions. Additionally, we introduced a second randomization strategy in the N- and C-cap as described^32, 49^ to allow also interaction of the capping repeats with the target. The libraries of DARPins with randomized and non-randomized N- and C- terminal caps, both containing randomized internal repeats and a stabilized C-cap, were mixed in a 1:1 stoichiometry to increase diversity. Successively enriched DARPin pools were cloned as intermediates in a ribosome display-specific vector^49^. Selections were performed over four rounds with decreasing target concentration and increasing washing steps to enrich for binders with slow off-rates and thus high affinities. The first round carried out the initial selection against myosin VI at low stringency. The second round included pre-panning with the undesired myosin VI isoforms 2 and 3 immobilized on magnetic beads, with the supernatant transferred to immobilized desired target myosin VI isoform 1. The third round included this pre-panning and the addition of non-biotinylated myosin VI isoform 1 to enrich for binders with slow off-rates. The fourth and final round included the pre-panning step and selection was performed with low stringency to collect all binders.

The final enriched pool was cloned as fusion construct with an N-terminal MRGS(H)_8_ tag and C-terminal FLAG tag via unique BamHI and HindIII sites into a bacterial pQE30 derivative vector containing lacI^q^ for expression control. After transformation of *E. coli* XL1-blue, 380 single DARPin clones were expressed in 96-well format and cells were lysed by addition of B-Per Direct detergent plus Lysozyme and Nuclease (Pierce). The resulting bacterial crude extracts of single DARPin clones were subsequently used in a Homogeneous Time Resolved Fluorescence (HTRF)-based screen to identify potential binders. The clone M6G4 that was selected for downstream applications was monoclonalized, by cutting the DARPin ORF, re-ligating it in fresh vector and retransformation. Binding of the FLAG-tagged DARPins to streptavidin-immobilized biotinylated ^His-Avi^myosin VI (aa 992-1031) was measured using FRET (donor: Streptavidin-Tb cryptate (610SATLB, Cisbio), acceptor: mAb anti FLAG M2-d2 (61FG2DLB, Cisbio). Further HTRF measurement against ‘No Target’ allowed for discrimination of myosin VI isoform 1-specific hits. Experiments were performed at room temperature in white 384-well Optiplate plates (PerkinElmer) using the Taglite assay buffer (Cisbio) at a final volume of 20 μl per well. FRET signals were recorded after an incubation time of 30 min using a Varioskan LUX Multimode Microplate (Thermo Fisher Scientific). HTRF ratios were obtained by dividing the acceptor signal (665 nm) by the donor signal (620 nm) and multiplying this value by 10,000 to derive the 665/620 ratio. The background signal was determined by using reagents in the absence of DARPins.

## Acknowledgements

We thank Ronald Wong and Kirill Petriukov for scientific discussions and Katharina Schlacher, Lorenza Penengo and Massimo Lopes for help with setting up the fiber assay. We thank Ron Hay, Simona Polo and Lorenza Penengo for sharing constructs or cell lines. We thank Thomas Reinberg, Sven Furler and Joana Marinho from HT-BSF UZH for their assistance in performing the ribosome display DARPin selection and screening. We thank the IMB Core Facilities for Proteomics, Flow Cytometry, Microscopy and Protein Production for technical support and reagents. This work was funded by the Deutsche Forschungsgemeinschaft (DFG, German Research Foundation) – Project-ID 393547839 – SFB 1361 awarded to H.D.U. and P.B., Project-ID BE 5342/2-1—FOR 2800 awarded to P.B. and Project-ID 408799149 awarded to H.P.W.

## Code availability

Custom code used for the preparation of volcano and GO term plots is available upon request.

## Supplemental information

**Fig. S1.**
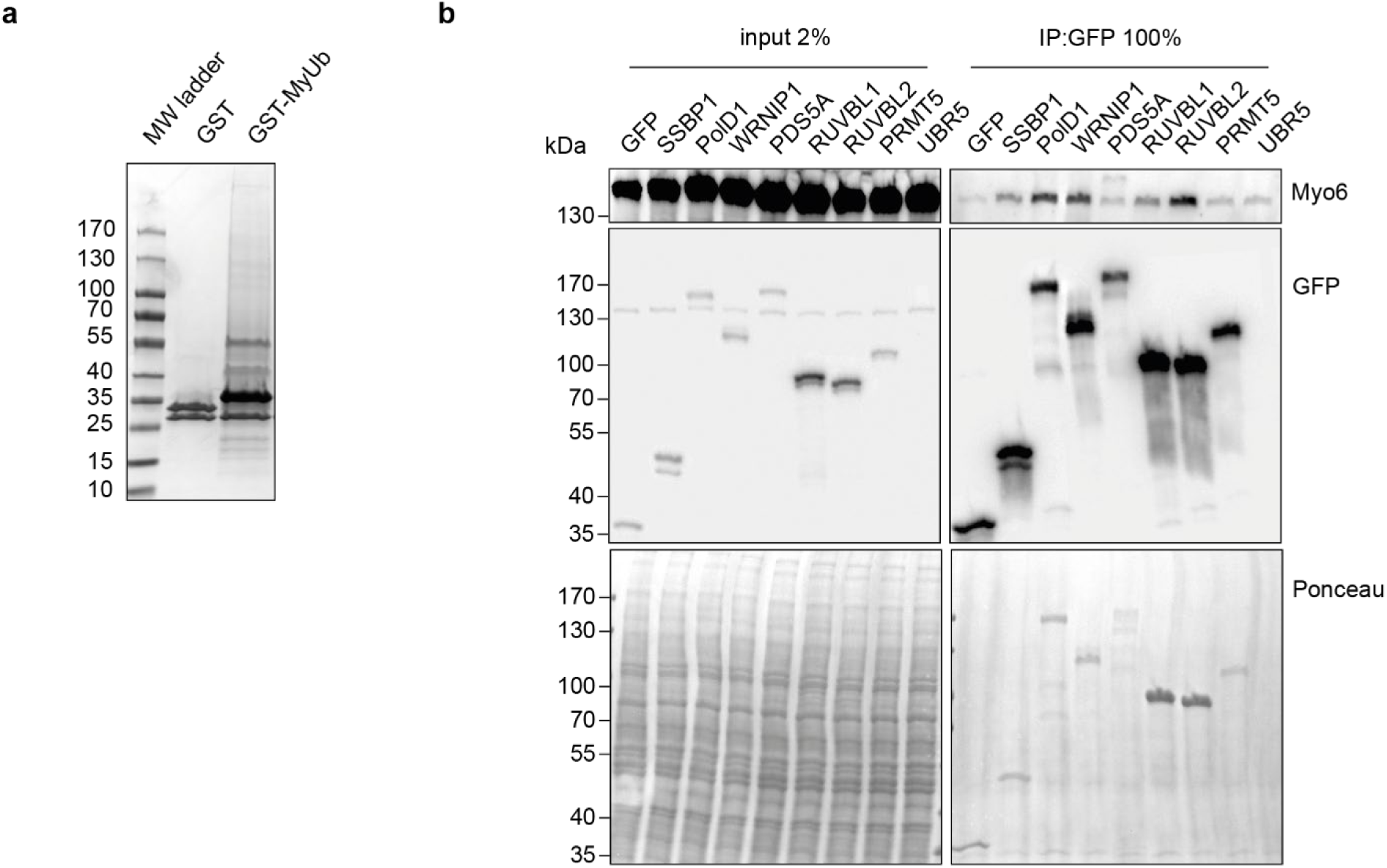
Identification of novel Myosin VI interactors. **a**, GST-pulldown assay using GST or GST-MyUb as bait and SILAC-labeled cellular lysates as prey. Samples were separated via SDS-PAGE and proteins were stained using a Colloidal Blue staining kit. **b**, Validation of novel interactors. HEK293T cells were transfected with GFP fusion constructs for the indicated proteins for 24 h. Immunoprecipitations (IPs) against GFP using GFP-trap beads (Chromotek) were performed, followed by western blotting with antibodies against myosin VI and GFP. Ponceau S staining was performed to control for equal loading.

**Fig. S2.**
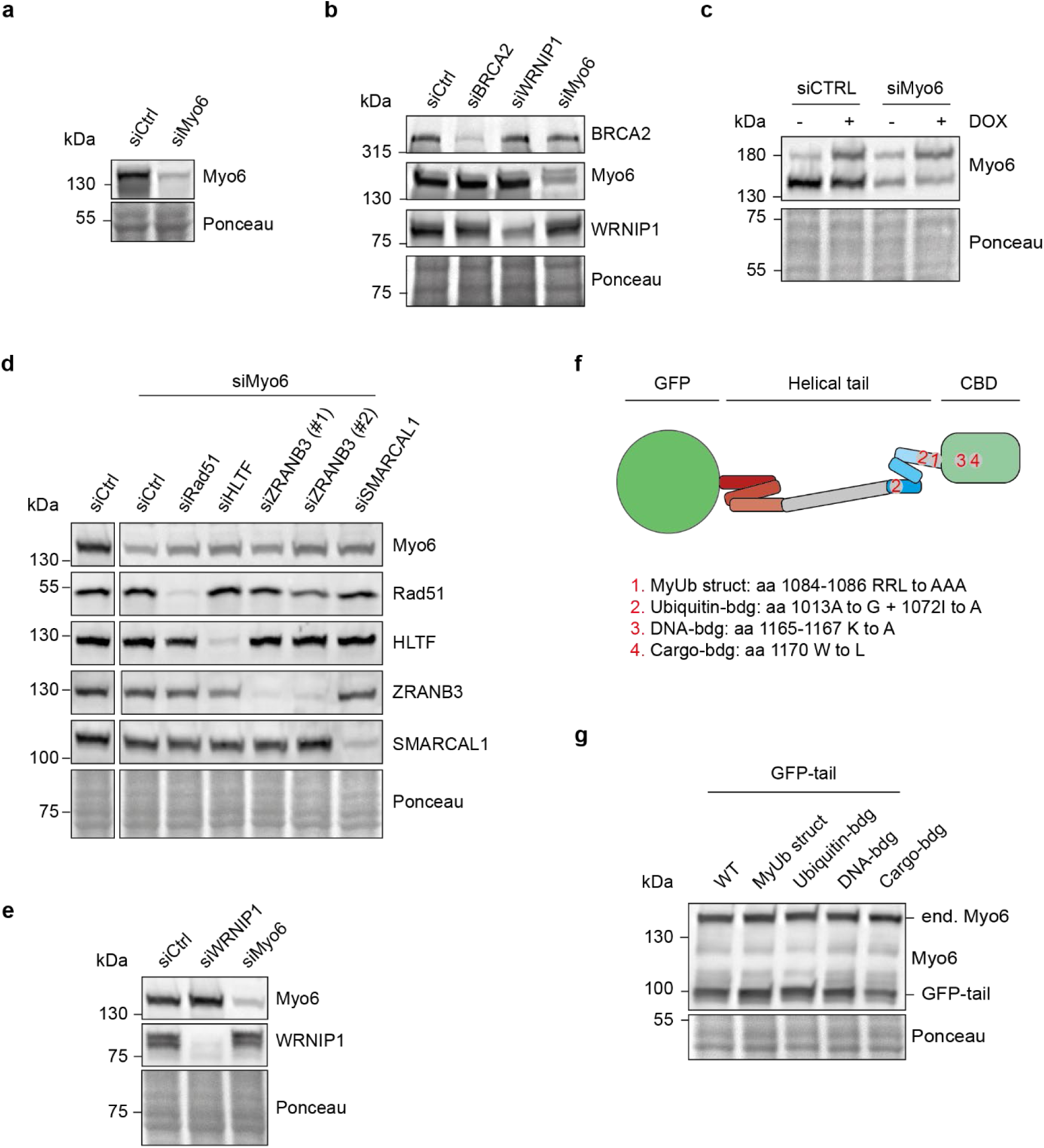
Knockdown and overexpression efficiencies in fiber assay experiments shown in. **Fig. 2** **a-e**,**g,** U2OS cells were transfected with the indicated siRNAs for 72 h or overexpression constructs for 48 h, followed by western blotting using the indicated antibodies. Ponceau S staining was performed to control for equal loading. Panel **a** relates to Fig. 2a, panel **b** relates to Fig. 2b, panel **c** relates to Fig. 2c, panel **d** relates to Fig. 2d, panel **e** relates to Fig. 2i and panel **g** relates to Fig. 2j. **f,** schematic representation of the respective GFP-tail mutants used in panel **g**

**Fig. S3.**
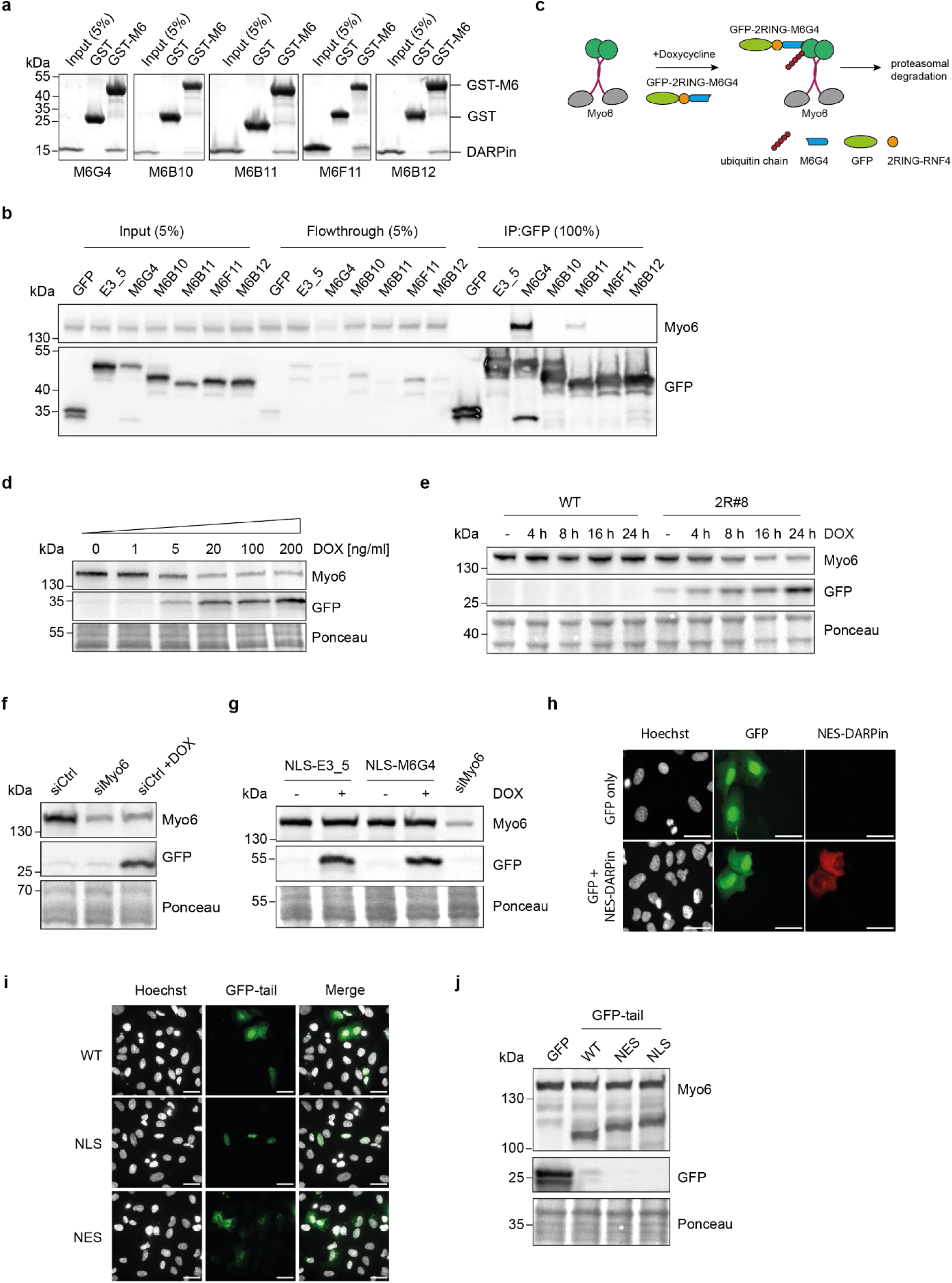
Establishment of DARPin-based tools and dominant-negative constructs to study contributions of cytoplasmic vs nuclear pools of myosin VI. **a**, DARPin screening via GST-pulldown assay. GST-pulldown assay using GST or GST-MyUb (labeled GST- M6) as bait and purified DARPin candidates as prey. Proteins were separated via SDS-PAGE and stained using Instant Blue. **b**, DARPin M6G4 depletes myosin VI from cellular lysates. HeLa cells were transfected with the indicated GFP-tagged DARPin constructs for 24 h. Immunoprecipitations (IPs) against GFP using GFP-trap beads (Chromotek) were performed, followed by western blotting with antibodies against myosin VI and GFP. **c**, Schematic representation of the DARPin-based myosin VI-degradation system adapted from Ibrahim et al.^1^. **d,e**, A U2OS Flp-In T-REx single cell clone harboring DOX-inducible GFP-M6G4-2RING fusion construct (2R#8) was treated with increasing concentrations of DOX for 24 h (**d**) or with 20 ng/ml DOX for the indicated times (**e**), followed by western blotting using antibodies against myosin VI and GFP. Ponceau S staining was performed to control for equal loading. **f,g,j**, Western blots to monitor knockdown efficiencies and expression levels for the fiber assays shown in Fig. 3. Panel f relates to Fig. 3c: U2OS Flp-In T-REx cells harboring DOX-inducible GFP-M6G4-2RING fusion construct (2R#8) were transfected with siRNAs for 72 h and treated with DOX for 24 h as indicated, followed by western blotting using antibodies against myosin VI and GFP. Panel g relates to Fig. 3e: U2OS Flp-In T-REx cells harboring DOX-inducible GFP-M6G4-NLS or GFP-E3_5-NLS (control) fusion constructs were transfected with siRNAs against myosin VI and treated with DOX as indicated, followed by western blotting using antibodies against myosin VI and GFP. Panel j relates to Fig. 3f: U2OS cells were transfected with GFP-tail constructs for 24 h as indicated, followed by western blotting using antibodies against myosin VI and GFP. Ponceau S staining was performed to control for equal loading. **h**, A GFP-selective NES-DARPin does not transport GFP from the nucleus to the cytoplasm. U2OS cells were co-transfected with GFP and mRuby-NES-tagged GFP-selective DARPin for 48 h as indicated. Nuclei were stained with Hoechst (white signal) (scale bar = 40 µm). **i**, Compartment-specific localization of tagged myosin VI-tails. U2OS cells were transfected with GFP or GFP-tail constructs as indicated. Nuclei were stained with Hoechst (white signal) (scale bar = 40 µm).

**Fig. S4.**
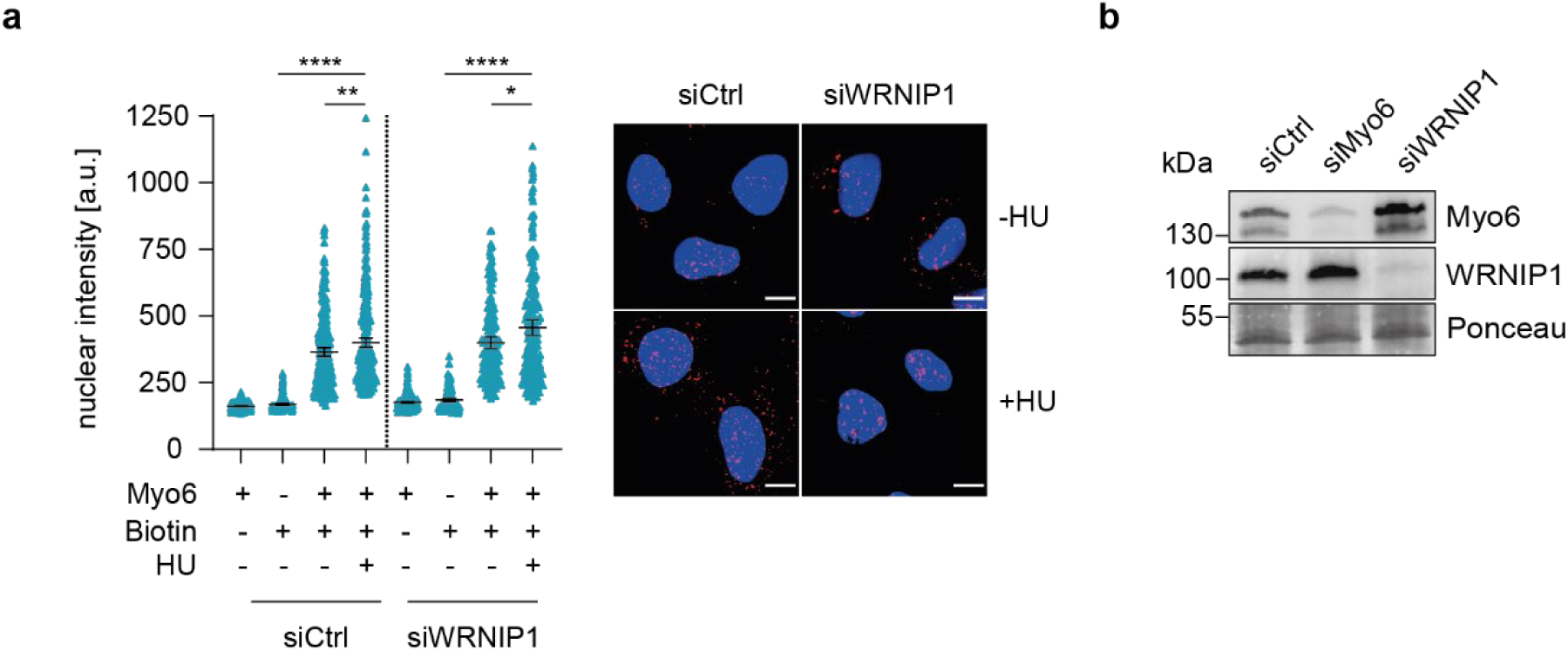
WRNIP1 depletion has no effect on the association of myosin VI with replication forks. **a**, U2OS cells were transfected with siRNAs for 72 h as indicated, followed by SIRF assay. Left: dot plots of PLA signal intensities with mean values -/+ 95 % confidence interval. Significance levels were calculated using the Mann-Whitney test from at least 100 nuclei per sample (ns, non-significant, ****: p<0.0001, ***: p<0.001, *: p<0.05). Right: representative images with Hoechst staining in blue and PLA signals in magenta, scale bar = 10 µm. A representative experiment from three independent replicates is shown. **b**, Western blot analysis to monitor knockdown efficiencies for SIRF assays shown in Fig. 4c and **S4a**.

**Supplementary Table 1.**
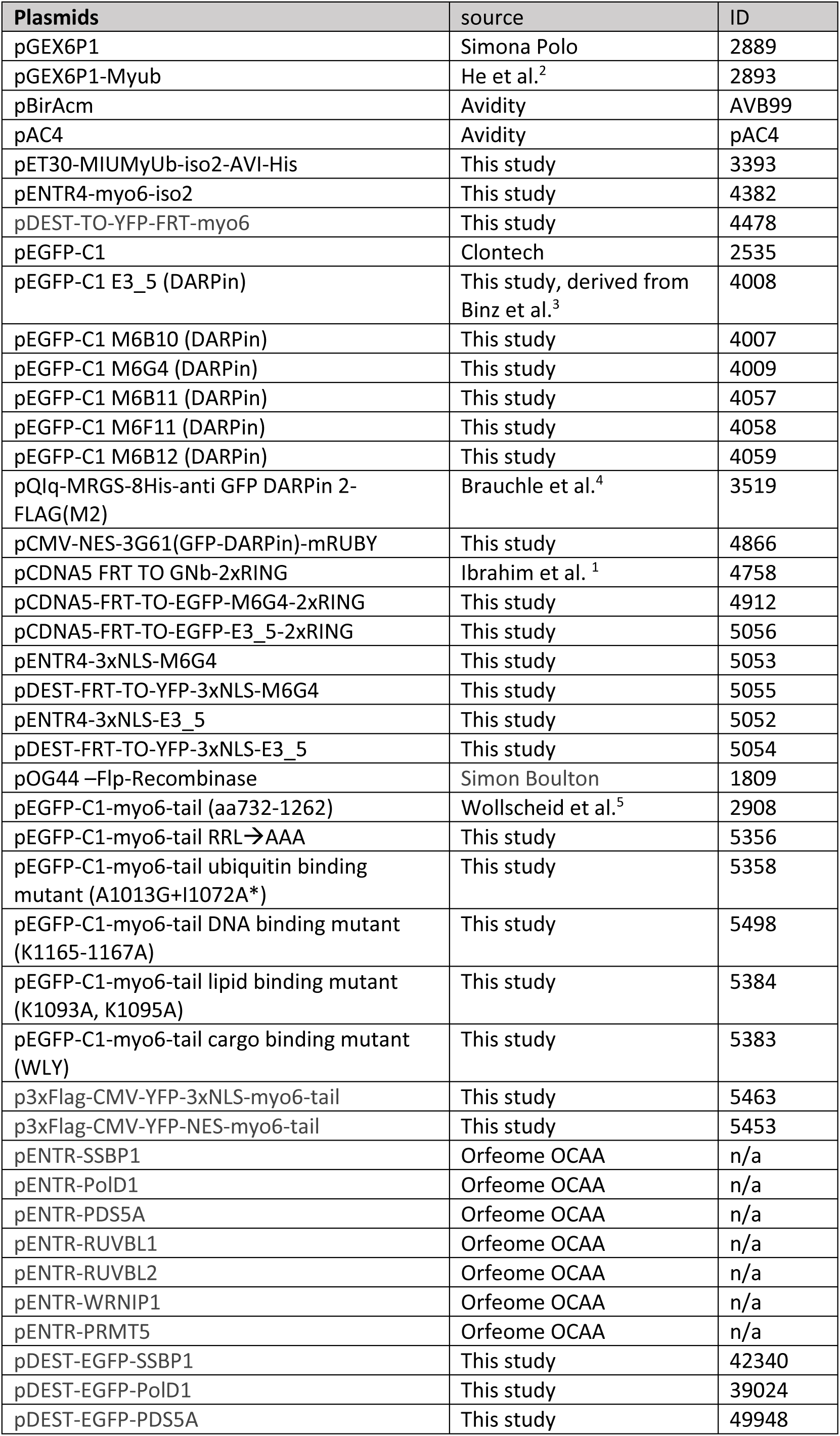
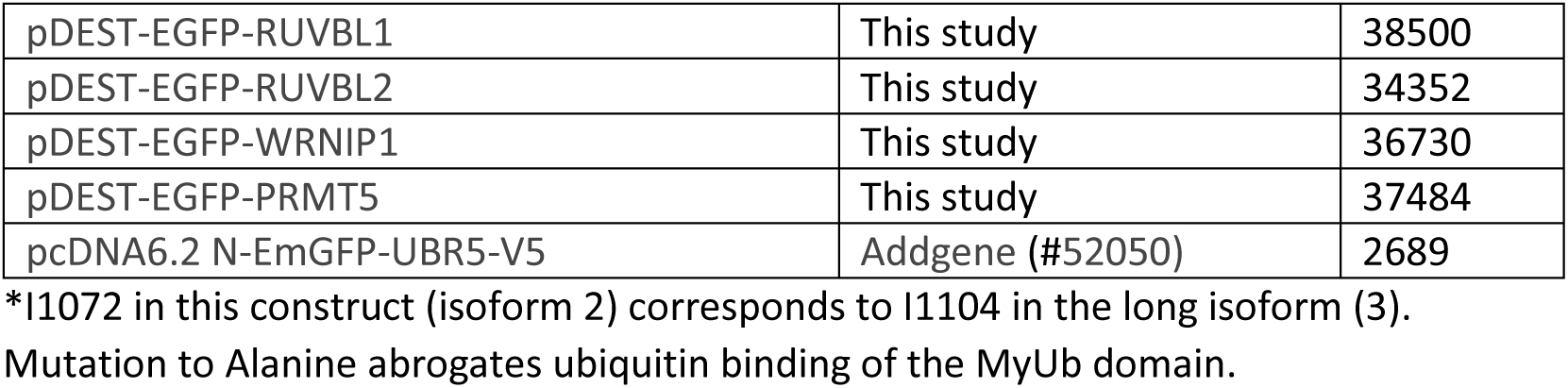
Plasmids used in this study

**Supplementary Table 2.**
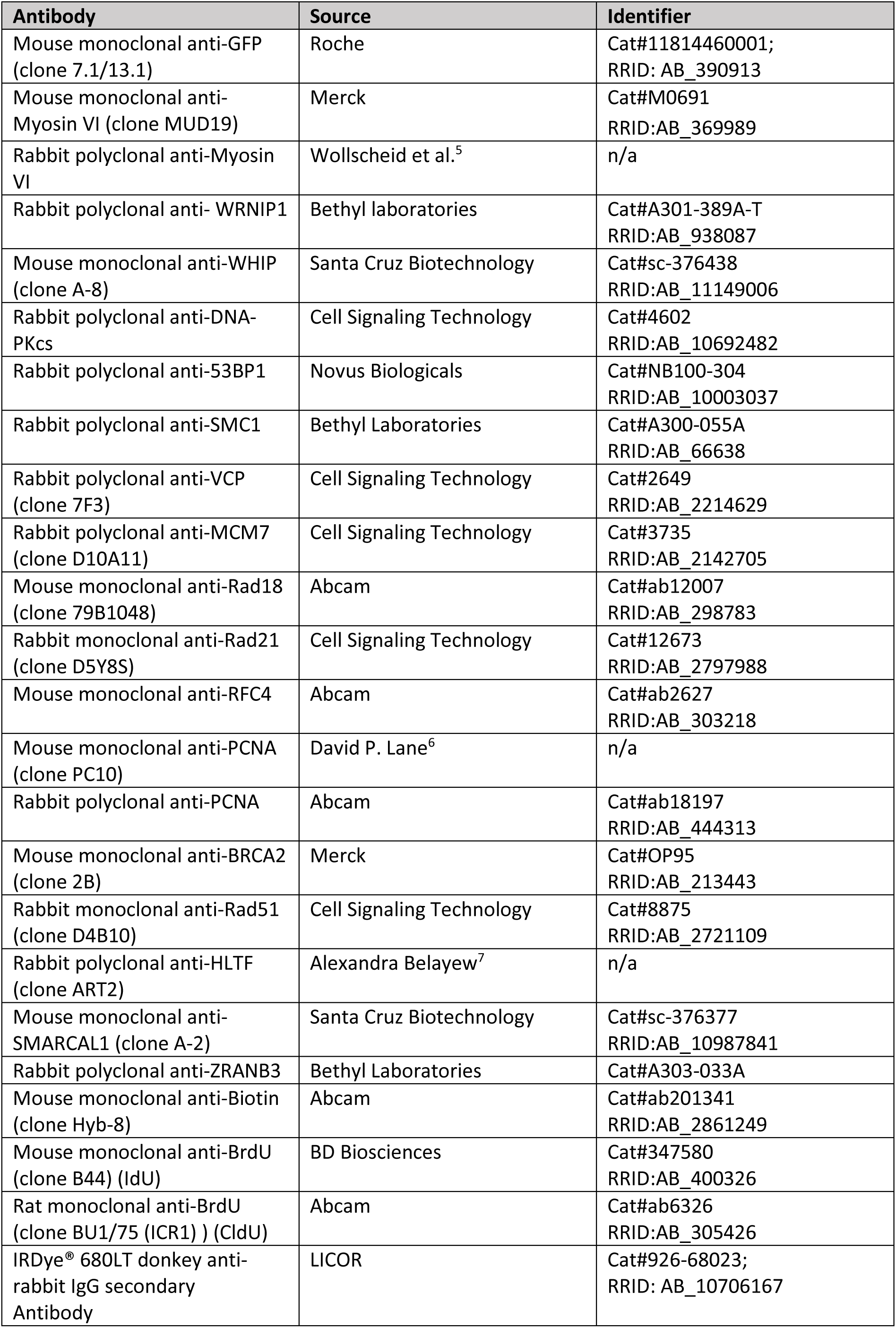
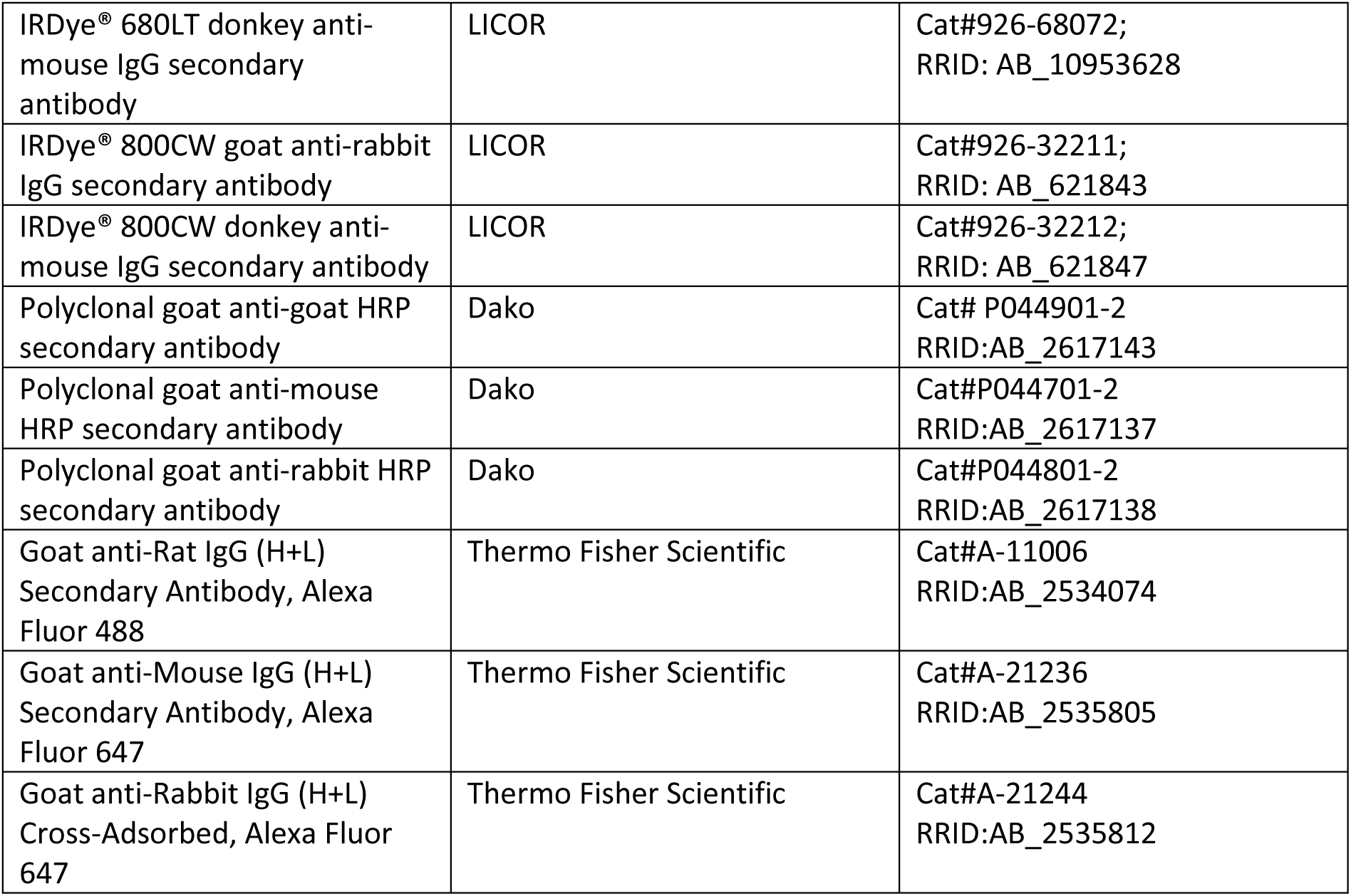
Antibodies used in this study

**Supplementary Table 3.** siRNAs used in this study

| siRNA | Source | Identifier |
| --- | --- | --- |
| Allstars Negative control siRNA | Qiagen | Cat#SI03650318 |
| Hs_MYO6_5 FlexiTube siRNA | Qiagen | Cat#SI03142692 |
| Hs_MYO6_7 FlexiTube siRNA | Qiagen | Cat#SI04243351 |
| Hs_MYO6_8 FlexiTube siRNA | Qiagen | Cat#SI04370737 |
| Hs_MYO6_10 FlexiTube siRNA | Qiagen | Cat#SI04998749 |
| FlexiTube GeneSolution GS56897 for WRNIP1 | Qiagen | Cat# GS56897 |
| FlexiTube GeneSolution GS675 for BRCA2 | Qiagen | Cat#GS675 |
| SiRNA Silencer Select [Hs] RAD51 | Thermo Fisher Scientific | Cat#s531930 |
| SiRNA Silencer Select [Hs] RAD51 | Thermo Fisher Scientific | Cat#s11734 |
| siRNA Silencer Select [Hs] HLTF | Thermo Fisher Scientific | Cat#s13137 |
| SiRNA Silencer Select [Hs] ZRANB3 | Thermo Fisher Scientific | Cat#s38488 |
| SiRNA Silencer Select [Hs] ZRANB3 | Thermo Fisher Scientific | Cat#s224929 |
| Hs_SMARCAL1_3 FlexiTube siRNA | Qiagen | Cat#SI00103194 |
| Hs_SMARCAL1_1 FlexiTube siRNA | Qiagen | Cat#SI00103180 |

**Supplementary Table 4.**
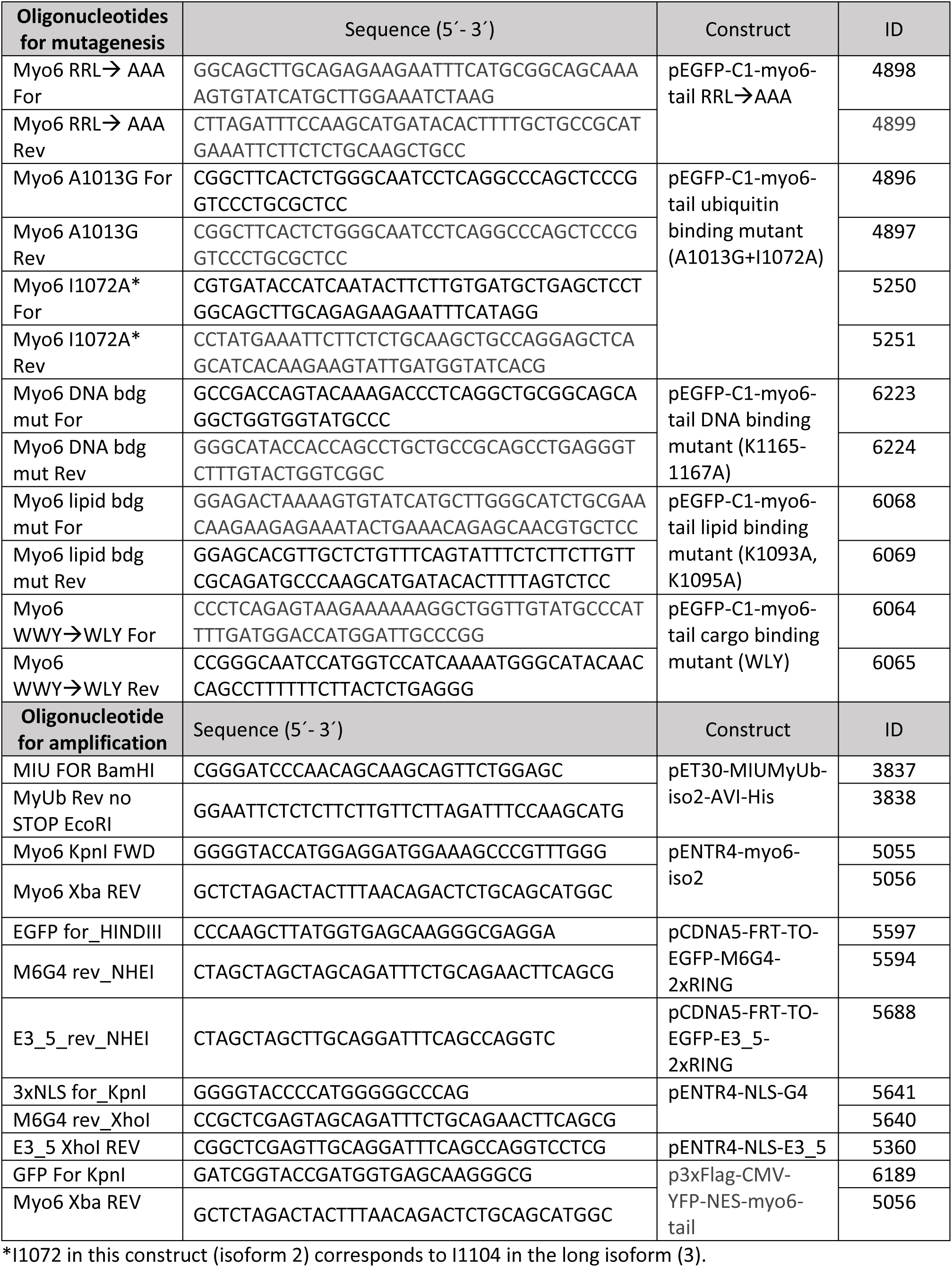
Oligonucleotides used in this study

**Supplementary Table 5.** Reagents used in this study

| **Chemical** | **Source** | **Identifier** |
| --- | --- | --- |
| Doxycycline hydrochloride | Merck | Cat#D3447 |
| 5-Chloro-2'-deoxyuridine | Merck | Cat#C6891 |
| 5-iodo-2'-desoxyuridin | Merck | Cat#I7125 |
| 5-ethynyl-2'-deoxyuridine | Merck | Cat#900584 |
| Azide-PEG3-biotin conjugate | Merck | Cat#762024 |
| Mirin | Merck | Cat#M9948 |
| DNA2 inhibitor C5 | AOBIOUS | Cat#AOB9082 |
| IPTG | Generon | Cat#GEN-S-02122 |
| Ni-NTA agarose | Qiagen | Cat#30250 |
| Imidazole | Merck | Cat#I2399 |
| Glutathione | Merck | Cat#G4251 |
| IPTG | Generon | Cat#GEN-S-02122 |
| Paraformaldehyde | Merck | Cat#P6148 |
| Formaldehyde solution, 36.5-38% in H<sub>2</sub>O | Merck | Cat#F8775 |
| Glycine | Merck | Cat#G7126 |
| Methanol | Thermo Fisher Scientific | Cat#15654570 |
| Acetic acid | Merck | Cat#A6283 |
| SIGMAFAST protease inhibitor cocktail | Merck | Cat#S8830 |
| cOmplete Protease Inhibitor Cocktail | Roche | Cat#5056489001 |
| Hygromycin B Gold | Invivogen | Cat#ant-hg-1 |
| Pfu Turbo DNA Polymerase | Agilent | Cat#600250 |
| DpnI | New England Biolabs | Cat#R0176L |
| Hydroxyurea | Merck | Cat#H8627 |
| Streptavidin Agarose Resin | Thermo Fisher Scientific | Cat#11846764 |
| Polyethyleneimine | Polysciences | Cat#23966-2 |
| Lipofectamine® 2000 | Thermo Fisher Scientific | Cat#11668019 |
| Lipofectamine® RNAiMAX | Thermo Fisher Scientific | Cat#13778150 |
| FuGENE® HD Transfection Reagent | Promega | Cat#E2311 |
| ProLong™ Diamond Antifade Mountant | Thermo Fisher Scientific | Cat#15205739 |
| DMEM, high glucose, pyruvate, no glutamine | Thermo Fisher Scientific | Cat#21969035 |
| Trypsin-EDTA (0.05%), phenol red | Thermo Fisher Scientific | Cat#25300054 |
| Penicillin-Streptomycin (10,000 U/mL) | Thermo Fisher Scientific | Cat#15140122 |
| L-Glutamine (200 mM) | Thermo Fisher Scientific |  |
| Blasticidin | Invivogen | Cat#ant-bl-1 |
| Triton X-100 | Merck | Cat#T9284 |
| Albumin (bovine serum albumin, BSA) | Merck | Cat#A7906 |
| Copper (II) sulfate | Merck | Cat#469130 |
| Sodium deoxycholate | Merck | Cat#D6750 |
| SDS, 20%, Sodium dodecyl sulfate | Merck | Cat#05030 |
| NuPAGE LDS Sample Buffer (4X) | Thermo Fisher Scientific | Cat#11559166 |
| GFP-Trap magnetic agarose beads | Chromotek | Cat#gtma-100 |
| Milk powder, skim milk | Merck | Cat#70166 |
| Tween-20 | Merck | Cat#P7949 |
| Amersham ECL Select Western Blotting Detection Reagent | Thermo Fisher Scientific | Cat#RPN2235 |
| Amersham ECL Prime Western Blotting Detection Reagent | Thermo Fisher Scientific | Cat#12994780 |
| Ponceau S | Merck | Cat#P3504 |
| Hoechst 33342 | Thermo Fisher Scientific | Cat#11534886 |
| Biotin | Merck | Cat#B4501 |
| PageRuler Prestained Protein Ladder | Thermo Fisher Scientific | Cat#11822124 |
| 4-15% Criterion™ TGX Stain-Free™ Protein Gel, 26 well, 15 µl | Bio-Rad Laboratories | Cat#567-8085 |
| Mini-PROTEAN TGX Stain Free Gels, 4-15%, 15-well | Bio-Rad Laboratories | Cat#456-8086 |
| SILAC light isotopes: L-Arg(0) and L-Lys(0) | Merck | Cat#A6969<br>Cat#L8662 |
| SILAC medium isotopes: L-Arg(6) and L-Lys(4) | Cambridge Isotope Laboratories | Cat#CLM-2265-H-1<br>Cat#DLM-2640-1 |
| Trypsin, MS-approved | Serva | Cat# 37286 |
| Glutathione Sepharose High Performance resin (GSH) | Cytiva | Cat#17527901 |
| **Critical commercial assays** |  |  |
| Duolink® In Situ Red Starter Kit Mouse/Rabbit | Merck | Cat#DUO92101-1KT |
| InstantBlue, 1L Protein Stain | Biozol | Cat#EXP-ISB01L |
| Trans-Blot® Turbo™ RTA Midi Nitrocellulose Transfer Kit | Bio Rad laboratories | Cat#1704271 |
| Gateway® LR Clonase® II enzyme mix | Thermo Fisher Scientific | Cat#10134992 |
| Colloidal Blue staining kit | Thermo Fisher Scientific | Cat#LC6025 |

